# Apoplastic effector candidates of a foliar forest pathogen trigger cell death in host and non-host plants

**DOI:** 10.1101/2021.08.06.455341

**Authors:** Lukas Hunziker, Mariana Tarallo, Keiko Gough, Melissa Guo, Cathy Hargreaves, Trevor S. Loo, Rebecca L. McDougal, Carl H. Mesarich, Rosie E. Bradshaw

**Author notes:** These authors contributed equally to this work (Lukas Hunziker, Mariana Tarallo). Centre for Crop and Disease Management, Curtin University, Bentley 6102, Perth, Australia (Lukas Hunziker).

## Abstract

Forests are under threat from pests, pathogens, and changing climate. One of the major forest pathogens worldwide is *Dothistroma septosporum*, which causes dothistroma needle blight (DNB) of pines. *D. septosporum* is a hemibiotrophic fungus related to well-studied Dothideomycete pathogens, such as *Cladosporium fulvum*. These pathogens use small secreted proteins, termed effectors, to facilitate the infection of their hosts. The same effectors, however, can be recognised by plants carrying corresponding immune receptors, resulting in resistance responses. Hence, effectors are increasingly being exploited to identify and select disease resistance in crop species. In gymnosperms, however, such research is scarce. We predicted and investigated apoplastic *D. septosporum* candidate effectors (DsCEs) using bioinformatics and plant-based experiments. We discovered secreted proteins that trigger cell death in the angiosperm *Nicotiana* spp., suggesting their recognition by immune receptors in non-host plants. In a first for foliar forest pathogens, we also developed a novel protein infiltration method to show that tissue-cultured pine shoots can respond with a cell death response to one of our DsCEs, as well as to a reference cell death-inducing protein. These results contribute to our understanding of forest pathogens and may ultimately provide clues to disease immunity in both commercial and natural forests.

## Introduction

Pests and pathogens are a persistent threat to plant health^1^, and this is likely to become worse with warming climate^2^. Forest trees, which provide important ecosystem functions and renewable resources, and have a role in mitigating global climate change^3^, are not exempt from this threat. Nonetheless, these functions are often undervalued^4^, and gymnosperm pathology research lags behind that of short-lived angiosperm crop plants.

Dothistroma needle blight (DNB) is one of the most destructive foliar diseases of pine trees. It has been reported in 76 countries, affecting 109 different host species^5^. DNB is caused by the Dothideomycete fungi *Dothistroma septosporum* and *D. pini*. These fungi colonise the needles, causing premature defoliation, reduced growth rates, and sometimes tree death^5,6^. *D. septosporum* accounts for most occurrences of DNB, particularly in North America and across the Southern Hemisphere, where it has devastated commercial plantations of *Pinus radiata*^7^. The increased incidence and severity of DNB seen over the last two decades has been attributed to both natural and anthropogenic causes, including climatic changes^8,9^. Existing DNB management strategies can counteract the disease in some cases, but huge losses are still seen in heavily affected regions, including some with native pine forests^5,7^.

To counteract any pathogen, it is important to understand how the pathogen achieves compatible interactions (infections) with its host plant(s). Of central importance to this compatibility is the host apoplast, where early contact between plant and pathogen cells is made^10^. For invading pathogens, the apoplast is a hostile environment, with constitutively produced plant molecules, such as secondary metabolites and hydrolytic enzymes, that impede growth^11^. In addition to these defences, the apoplast is monitored by cell surface-localised immune receptors, termed pattern-recognition receptors (PRRs). PRRs recognise foreign or ‘damaged self’ molecules, collectively called invasion patterns (IPs), to activate their innate immune system^12^. Following recognition by these PRRs, a series of immune responses of varying intensity are triggered, the strongest of which is the hypersensitive response (HR)^13^, which involves localised cell death and a burst of reactive oxygen species (ROS) that quickly halt the invading pathogen’s growth. Given this hostility, successful pathogens must neutralise the apoplastic environment to colonise their hosts and cause disease.

The ability to manipulate plant defences in the apoplast using secreted molecules is widely shared among microbes, including fungi^14^. These molecules, usually small proteins, are called effectors (virulence factors) and can evolve rapidly. However, following adaptation of the host, effectors that serve a virulence function for the pathogen may be recognised as IPs by host immune receptors and thus elicit resistance responses^10^. Immune receptors can also be located inside plant cells, where some effectors are delivered to perform their virulence functions. Intriguingly, some pathogens with necrotrophic stages in their lifecycles exploit plant cell death associated with IP recognition^15,16^. In these cases, the hosts’ attempts to prevent invasion instead favour pathogen growth by initiating cell necrosis, which provides nutrition for the pathogen.

Plant immune receptors that provide resistance against pathogens are encoded by resistance genes, and immune receptors that are hijacked by pathogens to provide susceptibility are encoded by susceptibility genes. The discovery of resistance and susceptibility genes in plants has averted yield losses for several crop species by enabling resistant cultivars to be selected^16^. Fuelled by ever-growing genome, transcriptome and proteome resources, the field of ‘effectoromics’, in which pathogen effectors are screened for those that elicit or suppress plant defence, has enabled the identification of hosts with resistance and susceptibility genes in a growing number of plant species^15–18^. Furthermore, these studies improve our understanding of host-pathogen interactions that will ultimately lead to new, more targeted, approaches to disease control.

Reports that investigate foliar pine pathogen effectors are scarce, with no efficient screening methods established involving the host species. *D. septosporum* is related to the well-studied tomato leaf mould pathogen, *Cladosporium fulvum*, with which it shares some functional effector genes^19–21^; both species are apoplast colonisers that do not use specialised infection structures like haustoria. The extracellular *D. septosporum* effector Ecp2-1 was recently suggested to be an avirulence factor which elicits defence responses in pine^22^. A recent study of *Pinus contorta* defence responses to *D. septosporum*^23^ identified upregulation of pine genes associated with IP-triggered immunity, and candidate immune receptor (*R*) genes showing positive selection, strongly suggesting the importance of *D. septosporum* effectors in triggering host defence. Here, we focused on candidate apoplastic effectors of *D. septosporum* that might be involved in a successful infection and/or triggering host defence. The specific aims of our study were to a) functionally characterise *D. septosporum* candidate effectors (DsCEs) based on recognition by conserved PRRs in non-host model plants, and b) develop and provide proof of concept for a new effector protein screening method in pines, employing vacuum infiltration-mediated delivery into small tissue-culture pine shoots.

## Results

### *D. septosporum* candidate effectors have sequence and structural similarity to fungal virulence factors

The *D. septosporum* genome has approximately 12,580 predicted genes, 397 of which encode putatively secreted proteins that are expressed during infection of *P. radiata* seedlings^19,24,25^. Using this resource, we identified apoplastic effector candidates of *D. septosporum* following the bioinformatic pipeline of Hunziker *et al* (2016)^25^ in conjunction with transcriptomics data to determine which genes were highly expressed and/or upregulated during infection of *P. radiata*^24^. We also used EffectorP 3.0^26^ and comparisons with the characterised apoplastic secretome of *C. fulvum*^21^ to generate a short list of 30 predicted apoplastic *D. septosporum* candidate effector proteins (DsCEs) (Fig. 1, Supplementary Table S1).

**Figure 1:**
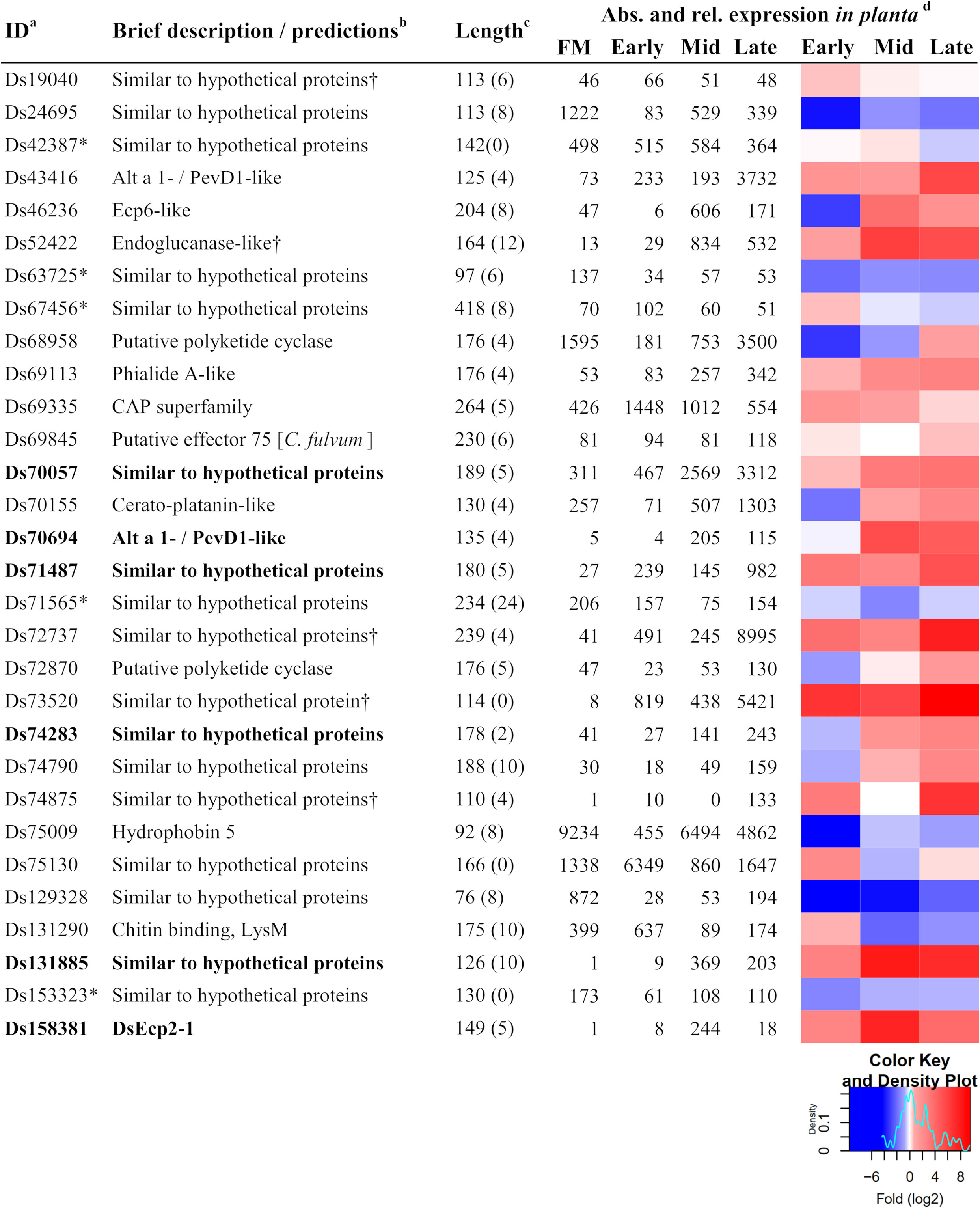
Cloned *Dothistroma septosporum* candidate effectors. Description and expression of the 30 shortlisted *D. septosporum* effector candidates used in this study. ^a^Protein ID numbers refer to those at JGI (https://mycocosm.jgi.doe.gov/Dotse1/Dotse1.home.html). An asterisk indicates a non-apoplastic localisation prediction according to EffectorP3.0. ^b^Function predictions were based on BLASTp (E<1e-05) and HHpred. A dagger indicates that no sequence homologues were found in other foliar pine pathogens for which predicted proteomes were available. ^c^Mature protein length (predicted N-terminal signal peptide removed) in amino acids; number of cysteine residues shown in brackets. ^d^*D. septosporum* gene expression in culture (FM; fungal mycelium) and *in planta* (*Pinus radiata*) at Early, Mid, Late infection stages ^24^. Left, Reads Per Million per Kilobase (RPMK); right, heatmap representing fold (log_2_) changes relative to FM expression.

To determine if the DsCEs were conserved in other fungi with available genomes, we used BLASTp to determine the top hits based on their predicted amino acid sequence similarity. All but five of the shortlisted DsCEs (Ds19040, Ds24625, Ds72737, Ds73520 and Ds74875) had homologues in at least 10 different fungal species, while nine were predicted to have conserved functional domains (Supplementary Table S2).

We then queried whether DsCE homologues are present in other fungi associated with pine needles, including seven pathogens and one endophyte for which predicted proteomes were available (Table S3). Three DsCEs (Ds19040, Ds72737 and Ds73520) appeared to be exclusive to *D. septosporum*, with no apparent homologues present in the included/queried pine needle-associated fungi. All other DsCEs, however, had a homologue in at least one other pine pathogen investigated (Supplementary Table S3), with more than half of these possessing a homologue in both a pathogenic species and the pine endophyte species *Lophodermium nitens*. Ds52422 was unique in having no homologues in the pathogenic species, but one homologue in *L. nitens* (Supplementary Table S3).

Recent studies showed that some effector protein families share structural similarity rather than sequence similarity with effectors from different fungi^27,28^. Thus the DsCEs were assessed for possible structural similarities to characterised proteins using HHpred. This analysis inferred structural relationships for five of the DsCEs that had no sequence-based functional annotations. Of these five, Ds43416 and Ds70694 showed structural similarity to Alt a 1 allergen / PevD1-like proteins, Ds52422 to endoglucanase proteins, and both Ds68958 and Ds72870 to polyketide cyclase proteins (Supplementary Table S1, Fig. 1). Notably, the gene encoding Ds43416 (Alt a 1 / PevD1-like) was one of the most highly expressed *and* up-regulated of the *DsCEs*, with a remarkably high level of expression at the late infection stage (Fig. 1, Supplementary Table S1). Among the other DsCEs assessed using HHpred, possible structural relationships were also identified with proteins that carry functional domains associated with fungal virulence, namely chitin binding (lectin), cerato-platanin (sub-group of endoglucanase-like proteins) and CAP family domains (Supplementary Table S1)^29–31^. In several cases, structural relationships inferred from HHpred corroborated functional annotations that were previously predicted for several DsCEs based on sequence similarity (Supplementary Table S1).

Taken together, the short list of 30 DsCE proteins (Fig. 1) included 13 with sequence and/or predicted structural similarity to fungal proteins with described functions, and 17 with only sequence similarity to yet undescribed proteins encoded in other fungal genomes (Fig. 1, Supplementary Tables S1 and S2). Twenty-five of the 30 DsCEs had at least four cysteine residues (Fig. 1). This suggests the presence of cysteine pairs forming disulphide bonds for stability in the hostile apoplast, as previously inferred for apoplastic effector proteins of *C. fulvum*^21^. Further, an overview of the *D. septosporum* reference genome, which is assembled to chromosome level^19^, showed no particular clustering of the 30 DsCE genes on small chromosomes or near repetitive elements (Supplementary Fig. S1, Supplementary Table S1). This corroborates earlier suggestions that, in contrast to several other Dothideomycetes, ‘pathogenesis-associated gene classes’ in *D. septosporum* were not enriched in these often-hyper-evolving regions^32^.

### Six candidate effectors of *D. septosporum* induced cell death in non-host plants

In previous work, the *D. septosporum* candidate effector DsEcp2-1 was shown to trigger cell death in *Nicotiana tabacum*, suggesting it might be recognised by an immune receptor in this plant species^22^. Similarly, small secreted proteins from the necrotrophic forest pathogen *Heterobasidion annosum* elicited cell death in *N. benthamiana*^33^. Thus, non-host angiosperm plants appear to recognise effector proteins from gymnosperm pathogens in a highly conserved fashion. To identify DsCEs that elicit cell death and are potentially recognised by non-host plant immune receptors, we therefore carried out *Agrobacterium tumefaciens*-mediated transient transformation assays (ATTAs) in *N. tabacum* and *N. benthamiana* with the 30 DsCEs listed in Fig. 1. In addition to DsEcp2-1, five other DsCEs induced cell death, indicative of a hypersensitive defence response (HR) (Fig. 2).

**Figure 2:**
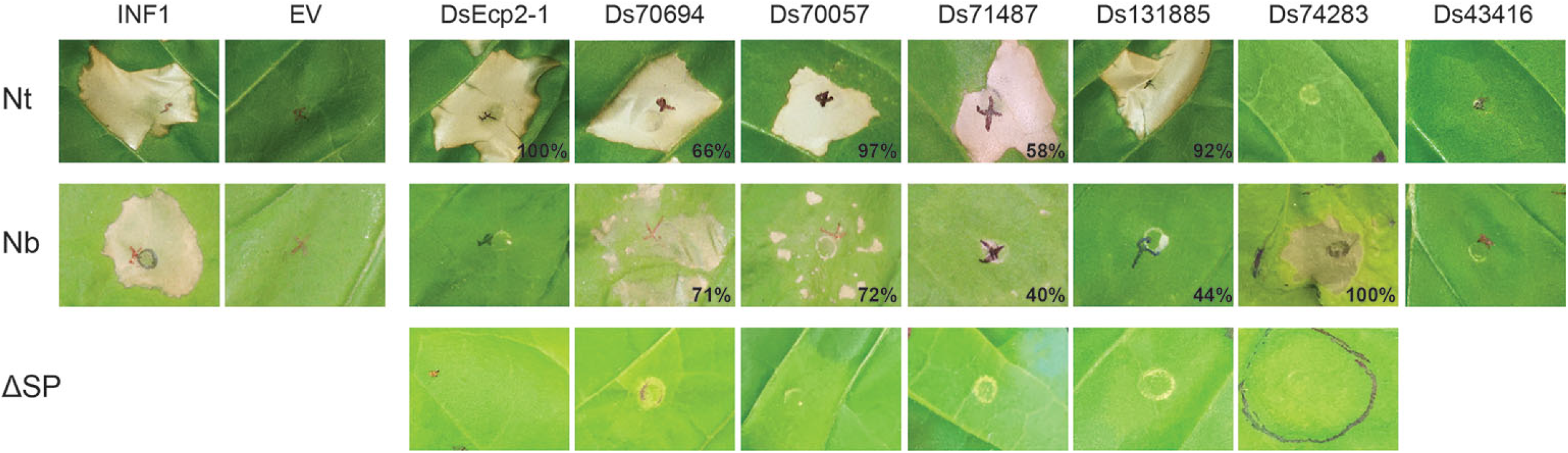
*Dothistroma septosporum* NZE10 candidate effectors (DsCEs) induce cell death responses in non-host plants. Genes encoding DsCEs, with a PR1α signal peptide for secretion to the plant apoplast, were introduced into *Nicotiana tabacum* (Nt; top row of images) and *N. benthamiana* (Nb; middle row of images) by agro-infiltration. Cell death responses were observed at 6 days post-infiltration. Representative images are shown, along with the percentages of all infiltrations where cell death occurred (n = 12 - 24 infiltration zones, from at least three independent experiments). Ds43416 is included as a representative of DsCEs that did not elicit a cell death response in either of the plant species tested. The bottom row of images shows the results of agro-infiltration of DsCE proteins lacking a PR1α signal peptide (ΔSP; no secretion into the apoplast) in *N. tabacum* (DsEcp2-1, Ds70694, Ds70057, Ds71487 and Ds131885) or *N. benthamiana* (Ds74283). INF1, *Phytophthora infestans* elicitin positive control; EV, empty vector negative control.

The DsCEs differed in their ability to consistently elicit a response in *N. tabacum*. Three (DsEcp2-1, Ds70057 and Ds131885) caused cell death in at least 90% of the infiltration spots within six days post-infiltration (dpi), while Ds70694 (66%) and Ds71487 (58%) were less consistent (Fig. 2). To gauge whether the culture density of infiltrated *A. tumefaciens* played a role in plant responses to the DsCEs, we trialled ATTAs at six optical densities (ODs; 0.05 - 1.0) with the three most consistent cell death-inducing DsCEs. Whilst DsCEs Ds70057 and Ds131885 triggered strong cell death across all ODs, DsEcp2-1 showed much weaker cell death at lower ODs (Supplementary Fig. S2), suggesting a difference in DsCE production *in planta*, or possibly a different type of interaction with plant targets.

The six DsCEs also differed in their ability to elicit cell death in the non-host plant *N. benthamiana*. Ds74283 was unique in eliciting consistent and strong cell death in *N. benthamiana*, despite showing no response in *N. tabacum*. Only two of the five effectors that induced cell death in *N. tabacum* (Ds70694 and Ds70057) also consistently induced cell death in *N. benthamiana*, although it was often patchy in the infiltrated zone (Fig. 2). Of the other DsCEs, Ds71487 and Ds131885 only induced weak cell death along the perimeter of infiltration zones in less than 50% of infiltrations into *N. benthamiana*, while DsEcp2-1 caused no cell death (Fig. 2).

To support the premise that the cell death responses were due to cell death elicitation by DsCEs in the apoplast, we cloned them for ATTAs without a secretion signal peptide. As expected, all of the non-secreted versions of these proteins failed to induce cell death (Fig. 2). The lack of cell death was not due to lack of DsCE production in the plant, but rather a lack of DsCE secretion to the apoplast, as each DsCE protein could be detected in infiltrated leaves by western blotting (Supplementary Fig. S3). Taken together, these results confirmed that the DsCEs need to be secreted into the apoplast to trigger cell death.

### *D. septosporum* cell death inducers are conserved among fungi

We queried the sequences of each DsCE protein that elicited cell death in *Nicotiana* spp. to further investigate their similarities to other fungal proteins. Some of these DsCEs are broadly conserved across different fungal classes, and even outside the Ascomycetes (Table 1, Table S4). Sequences similar to Ds70057 were the most numerous, with more than half of them found in Eurotiomycete species; at the other extreme, Ds70694 only had homologues in 14 Dothideomycete species. However, both Ds70694 and Ds70057 have multiple paralogues in some species, as well as in the *D. septosporum* genome, suggesting they are each members of a multi-gene family.

**Table 1:** BLASTp hits of *Dothistroma septosporum* cell death elicitors in fungal classes.

| Class<sup>a</sup> | Ds70057 | Ds70694 | Ds71487 | Ds74283 | Ds131885 |
| --- | --- | --- | --- | --- | --- |
| Dothideomycetes | 119 | 14 | 120 | 52 | 100 |
| Eurotiomycetes | 270 | 0 | 12 | 275 | 177 |
| Lecanoromycetes | 5 | 0 | 0 | 5 | 0 |
| Leotiomycetes | 30 | 0 | 38 | 26 | 14 |
| Orbiliomycetes | 0 | 0 | 0 | 1 | 0 |
| Pezizomycetes | 1 | 0 | 0 | 7 | 6 |
| Sordariomycetes | 77 | 0 | 157 | 10 | 70 |
| Taphrinomycetes | 0 | 0 | 2 | 0 | 1 |
| Xylonomycetes | 1 | 0 | 3 | 0 | 0 |
| Agaricomycetes | 4 | 0 | 0 | 0 | 0 |
| Pucciniomycetes | 0 | 0 | 0 | 0 | 12 |
| Ustilaginomycetes | 0 | 0 | 0 | 1 | 0 |
| Total hits | 507 | 14 | 332 | 377 | 380 |

<sup>a</sup> The numbers represent the number of species in these classes in which there was at least one BLASTp hit to the *D. septosporum* cell death elicitors ( $E < 1e-05$ ) from fungal species for which genomes were available at JGI (details in Supplementary Table S4). Ascomycetes are grouped above the line, Basidiomycetes below.

Among the cell-death elicitors, two were notable for their similarity to cell-death inducing proteins that have been described as PAMPs in other fungi. Based on reciprocal BLAST analyses, Ds71487 appears to be an orthologue of RcCDI1 from the barley pathogen *Rhynchosporium commune*^34^, with a pairwise identity of 41% and four conserved cysteine residues between the two proteins. Ds131885 is orthologous to VmE02, a cross-kingdom PAMP from the apple pathogen *Valsa mali* (Nie et al 2019), with 60% amino acid identity and ten conserved cysteine residues in both proteins.

### Development of a protein delivery method for pine tissue

Screening candidate effectors for their ability to elicit cell death in model plants is indicative of their potential functions. However, testing their functions in a pine host is critical for understanding their actual roles in disease. As a first step to achieving this, we developed a reliable, straight-forward methodology to deliver effector proteins into pine needles. Challenges included producing DsCE proteins of interest in sufficient purity and quantity, developing an effective protein delivery method, and identifying a positive control. Trials involved in the development of methods to address these challenges were discussed in detail by Hunziker (2018)^35^, and are briefly summarised here.

To produce the DsCE proteins of interest, we first trialled existing ATTA expression constructs and whole *N. benthamiana* leaves as expression systems for protein production. However, collection of apoplastic wash fluid (AWF), containing secreted DsCEs from infiltrated leaves, did not yield enough protein for replicated studies with pines. We then used heterologous protein expression and secretion in *Pichia pastoris*, which can be more easily up-scaled. Culture filtrates from *P. pastoris* transformed with an empty vector (negative control) elicited cell death in some *P. radiata* genotypes. Thus, to avoid this problematic background response, we purified histidine-tagged DsCEs from culture filtrates of *P. pastoris* using immobilised metal ion affinity (IMAC).

To deliver purified proteins into pine needles, we trialled several methods. Pine needles were not amenable to syringe infiltration, so vacuum infiltration, with pine tissue immersed in a DsCE protein solution, was used. After infiltration, detached needles, or groups of needles (fascicles), from pine seedlings quickly deteriorated, even when placed in moist conditions, so we chose different plant material for protein delivery assays. Here, clonal shoots of *P. radiata* produced by tissue culture without roots and maintained on agar medium, were employed^36^. The shoots were vacuum-infiltrated, then returned to the agar medium for 7–10 days. Vacuum infiltration of whole shoots with a neutral red dye solution suggested good uptake efficiency. A schematic of the final method is shown in Fig. 3.

**Figure 3:**
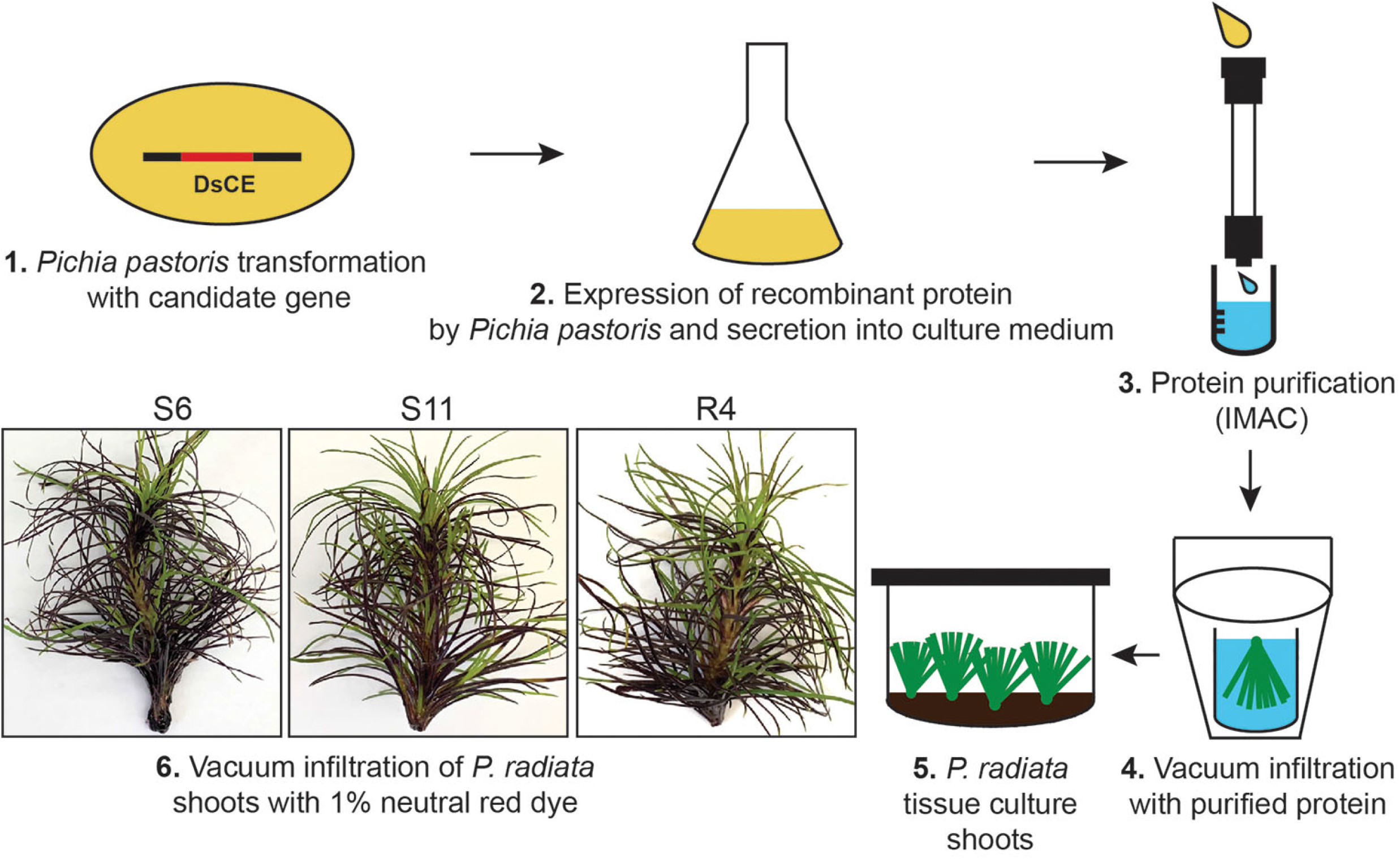
Overview of the pine shoot vacuum infiltration method developed in this study. *Dothistroma septosporum* candidate effector Ds70057 and positive control *Botrytis cinerea* effector BcSSP2 proteins were produced through heterologous expression and secretion by *Pichia pastoris* (1,2). The proteins were purified by immobilized metal affinity chromatography (IMAC) (3) then vacuum-infiltrated into clonal microshoots of *Pinus radiata* (4). Infiltrated shoots were returned to LPch media for up to 7 days after infiltration (5). 1% neutral red was infiltrated to assess the efficiency of vacuum infiltration (6) in the different pine genotypes Sus6 (S6), Sus11 (S11) and Res4 (R4).

A positive control that elicits cell death when infiltrated into clonal shoots of *P. radiata* was required to confirm the efficacy of the new method. An effector protein from the broad host-range pathogen *Botrytis cinerea*, BcSSP2, was previously shown to induce strong cell death in *N. benthamiana* as well as in other plants^37^, so was tested in this study. Consistent with the previous study, vacuum infiltration of purified BcSSP2 protein produced by *P. pastoris* elicited cell death in three different genotypes of pine shoots within three days, providing a positive control. Importantly, the negative control (protein purification elution buffer) did not elicit a response (Fig. 4).

**Figure 4:**
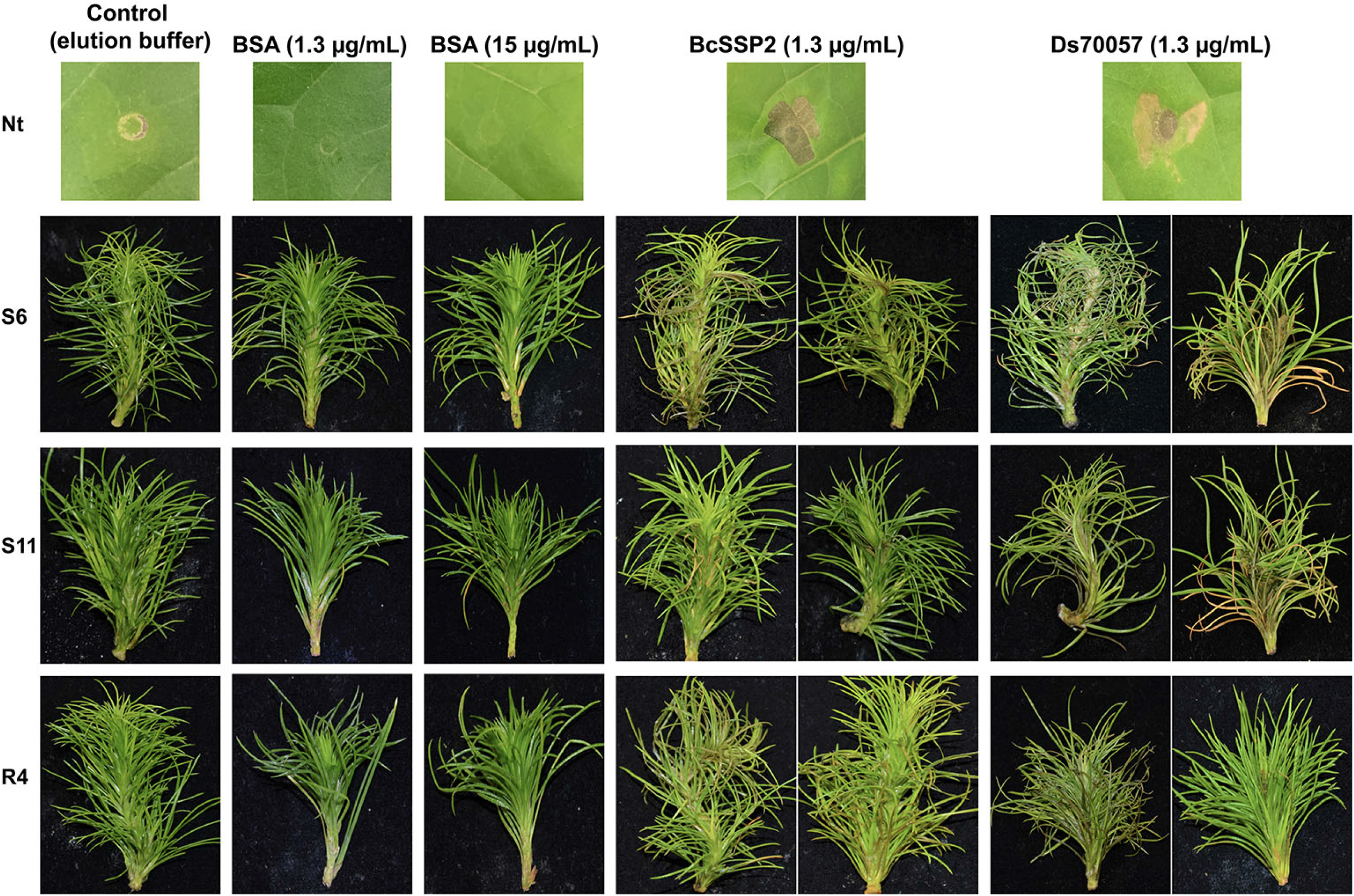
The *Dothistroma septosporum* candidate effector (DsCE) Ds70057 induces cell death in *Pinus radiata* shoots following vacuum infiltration. Whole shoots of *P. radiata* genotypes that are susceptible (S6 and S11) or tolerant (R4) to *D. septosporum* infection were infiltrated with purified *Botrytis cinerea* SSP2 (positive control) or DsCE Ds70057 protein produced by heterologous expression in *Pichia pastoris*. Negative controls were elution buffer only and solutions of bovine serum albumin (BSA). Representative photos (from 18-24 pine shoots for each treatment) were taken 7 days after infiltration. The top panel shows infiltrations of the respective solutions into *Nicotiana tabacum* (Nt) for reference.

### Infiltration of a DsCE protein into pine shoots induces cell death

With a novel protein delivery method established using pine shoots, we tested one of the DsCE proteins, Ds70057, to investigate its capacity to trigger cell death in pine like in *N. tabacum*. Ds70057 was selected because it was highly expressed at the ‘mid’ and ‘late’ (necrotrophic) stages when lesions are formed by *D. septosporum* in *P. radiata*^24^, and as such was seen as a candidate for playing a role in the switch from biotrophic to necrotrophic growth.

*Pinus radiata* shoots were infiltrated with the Ds70057 purified protein solution (1.3 μg/ml), alongside controls. As expected, the negative controls (elution buffer, bovine serum albumin) did not cause any damage or visible stress responses (Fig. 4). In contrast, Ds70057 consistently induced cell death within seven days across all three *P. radiata* genotypes tested (S6, S11 and R4) (Fig. 4, Supplementary Fig. S4), with these responses near-identical to the positive control BcSSP2. This finding suggests that Ds70057 might be a virulence factor in the necrotrophic stage of dothistroma needle blight that helps to destroy needle tissue. More importantly, we showed that *P. radiata* needles can be screened for responses to potential effectors.

## Discussion

Dothistroma needle blight (DNB) remains a threat to both commercial and native pine forests worldwide, despite decades of research into this disease and its management^5,7^. Although screening with pathogen effectors based on genome data (effectoromics) has fast-tracked identification of resistant angiosperm crop varieties with corresponding immune receptors, it is not known whether a similar effectoromics-based approach could be applied to improve forest health given the longer life-cycles of forest trees, and the complexity of factors affecting disease resistance^38^. However, a recent transcriptomics study of *Pinus contorta* responses to infection with *D. septosporum* showed up-regulation of pine genes involved in Ca^2+^ and MAPK defence signalling pathways as well as pathogen-specific defence responses^23^. Signatures of selection were identified in candidate immune receptors (*R* genes), suggesting adaptation to the pathogen and the importance of specific interactions between pathogen effectors and host immune receptors in pine defence to *D. septosporum*.

Here we used an effectoromics screening approach in two non-hosts to explore the effector arsenal of *D. septosporum*. Of 30 small secreted *D. septosporum* candidate effector (DsCE) proteins that were expressed by transient transformation in *N. tabacum* or *N. benthamiana*, six induced cell death. Four of these six proteins are very highly conserved across a broad range of fungal taxa, suggesting some commonality in angiosperm and gymnosperm effectors, as also seen in some *Phytophthora* pathogens^39^. Most effectoromics studies include screening on host as well as non-host plants but, to the best of our knowledge, there were no methodologies in place to enable effector protein screening on pines or indeed any other gymnosperms. Thus, we developed a novel method that involves infiltration of purified effector proteins into clonal pine shoots. We validated the system using protein of a highly expressed DsCE, Ds70057, that elicited cell death in *N. tabacum*, and found that it also triggered cell death across three independent clonal genotypes of *P. radiata*, suggesting that an effectoromics screen for host resistance may be possible even in gymnosperm trees.

We built on a prior study^25^ to identify and refine a small set of secreted proteins that are likely to play a role as effectors in the infection of *P. radiata* by *D. septosporum*. A major selection criterion for these proteins was high expression and/or up-regulation during the infection ^24^. We mainly adhered to size and cysteine content thresholds (≤300 aa, ≥4 Cys) used in studies of angiosperm pathogens^40^ and used the EffectorP 3.0^26^ tool to help classify the DsCE candidates.

Functional predictions and screens for homology usually rely on comparisons of primary amino acid sequences using BLASTp. In our study, we also used HHpred to infer possible structural relationships between the 30 DsCEs and proteins with characterized tertiary structures. Notably, HHpred predictions largely supported the BLASTp results, but also uncovered potential functions for five additional DsCEs. One of these five, Ds70694, induced cell death in *Nicotiana* spp. and was predicted to have structural similarity to proteins with the major human allergen Alt a 1 fold^41^, with the top HHpred hit to the PAMP-like effector protein PevD1 from *Verticillium dahliae*^42^. Another protein with an Alt a1 fold, the *Magnaporthe oryzae* HR-inducing protein 1 (MoHrip1), is highly induced during the infection of its rice host and has been implicated in promoting disease^43^. Notably, it has been shown that, like Ds70694, both PevD1 and MoHrip1 can trigger cell death in *Nicotiana* species^42–44^. This is despite the fact that *Nicotiana* species are not hosts for *D. septosporum*, *V. dahliae* or *M. oryzae*, suggesting that all three proteins share a conserved function in promoting cell death and/or they possess a conserved epitope that is recognised as a PAMP by these plant species.

Another of the cell-death inducing DsCEs, Ds131885, is an orthologue of VmE02, a cross-kingdom PAMP from the necrotrophic apple pathogen *Valsa mali*^45^ that also elicits cell death in *Nicotiana* spp.. *VmE02* mutants had reduced numbers of pycnidia, suggesting the protein may be involved in pathogen conidiation. A similar function for Ds131885 would concur with its upregulated expression at the mid and late stages of infection. Recently a receptor-like protein (RLP) in *N. benthamiana* was identified that mediates VmE02-triggered immunity to fungal and oomycete pathogens^46^.

Ds71487 is homologous to a *Rhynchosporium commune* protein, RcCDI1, which induced cell death in *N. benthamiana*, *N. sylvestris*, and two other Solanaceae species, but not in its host barley^34^. RcCDI1 homologues in three other pathogens also induced cell death in *N. benthamiana*, and the authors concluded that the protein is a PAMP^34^. Given this context, Ds71487 inducing cell death in *N. tabacum* is not surprising and this response might also be found in other plants. However, it is unclear why this protein only triggered a weak response in *N. benthamiana*.

The *D. septosporum* effector protein DsEcp2-1 was already known to elicit cell death in *N. tabacum* from a previous study^22^. In that work, *DsEcp2-1*-deficient mutants showed increased virulence on *P. radiata* compared to an *Ecp2-1*-containing wild-type strain, suggesting a possible avirulence role for DsEcp2-1 in eliciting a defence response by the plant. This suggestion supports the concept that *D. septosporum* effectors could, through adaptation of the host, become a warning signal (avirulence effector) similar to those of biotrophic pathogens^47,48^.

The remaining DsCEs that caused cell death in non-host plants in our study were not affiliated with any known functions, but were highly conserved across a broad range of fungal taxa (Table 1). Effectors with conserved core functions shared among plant pathogens could be important in elicitation of plant resistance as they are likely to be essential for the microbes’ survival. Mutation or loss of essential conserved pathogen effector genes to escape recognition by a specific plant host immune receptor would most likely lower pathogen viability. It is possible that highly conserved ‘core’ effectors may have been overrepresented in our results because we screened gymnosperm pathogen effectors in angiosperm non-host plants. However, our results strongly affirm an earlier study that suggests conservation of core effectors between pathogens of angiosperms and gymnosperms^39^. The matching cell-death responses of pine and *Nicotiana* sp. to infiltration with Ds70057 and the control *B. cinerea* SSP2 proteins in our study also suggest similarities in the physiological responses of these disparate plant groups. Similarly, Chen et al. (2015)^49^ found that infiltration of the cerato-platanin protein (CPP) HaCPL2 from *Heterobasidion annosum* induced cell death in both the host *Pinus sylvestris* when applied to roots *in vitro* as well as in *N. tabacum* leaves.

Effectors can sometimes have opposing roles depending on the host context. Whilst they are mainly exploited in effectoromics studies on the basis of a role in avirulence, in which their recognition by a specific plant immune receptor elicits defence, they can have important roles in virulence that may be indispensable for fitness, causing conflicting selection pressures. At the same time, the induction of cell death by the plant host as part of its defence mechanism is beneficial for necrotrophic pathogens, such as *Parastagonospora nodorum* that causes Septoria blotch of wheat^15^. The *P. nodorum*-wheat interaction is regarded as a model pathosystem for understanding the ‘inverse gene-for-gene’ relationship between effectors and their corresponding plant targets, which are termed susceptibility factors. For both biotrophic and ‘inverse’ necrotrophic interactions, germplasm can be screened with effector proteins for a resistant or susceptible reaction, respectively, and accordingly selected or deselected for breeding^16^. However, the situation with hemibiotrophs, such as that of *D. septosporum*, is more complex and requires accurate understanding of the disease.

The DsCE Ds70057 is of interest due to its very high expression and up-regulation in the mid and late (necrotrophic) stages of disease caused by *D. septosporum* on *P. radiata*^24^ and its ability to elicit cell death in three genotypes of *P. radiata* as well as in *N. tabacum*. We suggest it may be a necrotrophic effector triggering cell death that assists the pathogen by eliciting the destruction of needle tissue. This action would augment that of dothistromin which is required for disease lesion expansion, but not for initial elicitation of cell death^50^. Further studies with Ds70057, including construction and analysis of knockout mutants, will help to determine if Ds70057 is acting as a necrotrophic effector, in which case we would expect mutants to have decreased virulence.

An aim of our work was to develop a method to screen putative effectors of conifer pathogens directly in their host using a protein infiltration approach, with the long-term goal of identifying corresponding immune receptors for the effectors, and thus assisting breeding for disease resistance. There were several challenges associated with protein delivery into pines. In an earlier study on *P. sylvestris*, roots took up purified protein from a root pathogen via filter paper strips, resulting in root cell death^49^. By submerging pine shoots into solutions containing purified candidate effector proteins, and applying a vacuum pressure, we were able to develop a robust and reliable infiltration process that did not harm the pine needle tissue. We identified a positive control that can be used as a reference in future studies and showed that a secreted protein of *D. septosporum* induced cell death symptoms in pine shoots as well as *Nicotiana* leaves.

This novel method of effector delivery to gymnosperm tissue would be well suited for medium-scale screening of multiple pine genotypes with a small number of effector proteins. If the effector-screening process was to be scaled up for high-throughput commercial use with a large number of effector proteins, the time used to grow and prepare fresh purified effector proteins in the *P. pastoris* system as used here could be a limiting factor. An alternative delivery method might be through *Agrobacterium*-based transformation of pines, in which the effector genes rather than the proteins are delivered to the host plant. *Agrobacterium*-based transformation of *P. radiata* and other *Pinus* spp. has been achieved^51,52^, but it is a slow, technically challenging process. Despite this, future efforts to establish reliable transient transformation of pines may be worthwhile.

This study has broad implications for our understanding of fungal pathogens of gymnosperm trees and how these species interact with each other at the molecular level. Importantly, the high level of conservation of effectors across fungal taxa, and conservation of plant responses across angiosperms and gymnosperms suggests that much of what the scientific community has learned about plant-pathogen interactions involving angiosperm systems may also be relevant to gymnosperms. A similar conclusion was reached by studying pine responses to *D. septosporum* infection^23^. There also appear to be common ‘core’ interactions that could be exploited to give broad host-range resistance. Another implication of our work relates to the pine screening method which could be adapted for use with other gymnosperm tree species and other plant hosts that are not amenable to the usual screening methods. Finally, this type of screening method has the potential, along with expression data and pathogen gene knockout studies, to identify necrotrophic effectors that could in turn identify susceptibility gene targets, leading to durable resistance.

## Methods

### Bacterial strains and plants

*Escherichia coli* DH5α was used for gene cloning and plasmid propagation. *A. tumefaciens* GV3101^53^ was used, in conjunction with *N. tabacum* Wisconsin 38 and *N. benthamiana*, for ATTA experiments. *P. pastoris* GS115 was used for heterologous protein expression in liquid culture. *P. radiata* clonal shoots without roots, grown from embryo cotyledon tissue under sterile conditions on LPch agar^36^, were used for protein infiltration experiments.

### DsCE identification and homology searches

Genes encoding conserved candidate secreted apoplastic effector proteins were identified in the *D. septosporum* genome ^19^, based on a bioinformatic pipeline published previously^25^. A shortlist of 30 putative apoplastic effector proteins was determined based on predictions that they were secreted ^25^, apoplastic - as determined using EffectorP3.0^26^ and expressed during the infection of *Pinus radiata* seedlings^24^.

BLASTp was used to identify the top homologs of the 30 DsCE proteins in the National Center for Biotechnology Information (NCBI) and Joint Genome Institute (JGI) databases, with an Expected (E)-value threshold of 1E-02. This included a specific query against the foliar pine pathogens *Fusarium circinatum* (PRJNA565749), *Lecanosticta acicola* (PRJNA212329), *Cronartium ribicola* (PRJNA190829), *Pseudocercospora pini-densiflorae* (PRJNA212512), *Cronartium quercuum* f. sp. *fusiforme* (PRJNA67371), *Elytroderma deformans* (PRJNA537175), *Gremmeniella abietina* (PRJNA347218) and *Cyclaneusma minus* (R. McDougal, Scion, unpublished), as well as the pine endophyte *Lophodermium nitens* (PRJNA335148) (Supplementary Table S3). BLASTp was also used to determine the level of conservation for each of the cell death-eliciting DsCEs across fungal species in JGI, with an E-value threshold of 1E-05. Here, only one isolate per species was examined, with all paralogs from each species retained. Conserved functional domains in the DsCE proteins were identified using the Conserved Domain Database (CDD) in NCBI.

HHpred^54^ was used to infer possible structural relationships between the 30 DsCEs and proteins of characterized tertiary structure present in the RCSB (Research Collaboratory for Structural Bioinformatics) protein databank. Here, only the top hit was retained. Structural relationships were deemed to be significant if they had a probability score of 95%, an E-value score <1E-03, and an overall score of >50. Despite not meeting these criteria, Ds43416 and Ds70694 were also deemed to have significant structural relationships with homologous proteins of *C. fulvum* through additional support from previous structural predictions^21^.

### Generation of DsCE expression vectors for *Agrobacterium tumefaciens*-mediated transient transformation assays (ATTA)

DsCE ATTA expression vectors were generated using a method described previously^22^. Briefly, *DsCE* genes, without their native signal peptide sequence, were PCR-amplified from *D. septosporum* NZE10 genomic DNA or cDNA (derived from *in vitro* or *in planta*-grown fungus^24^) using custom primers (Supplementary Table S5; Integrated DNA Technologies) with *Bsa*I recognition sites and overhangs required for modular assembly using the Golden Gate approach^55^. PCR amplicons were then directly used as entry modules for cloning into the ATTA expression vector pICH86988, which contains a CaMV 35S promoter and octopine synthase terminator flanking *Bsa*I insertion sites^56^, using Golden Gate assembly, along with an entry module encoding an N-terminal PR1α signal peptide for secretion into the apoplast and an N-terminal 3 X FLAG tag for detection by western blotting. Cell death-eliciting DsCEs were also assembled into expression vectors as above without a PR1α signal peptide. DsCE ATTA expression vectors were then transformed into *E. coli*, and inserts confirmed by sequencing. Finally, the DsCE ATTA expression vectors, along with the INF1 ATTA expression vector (an extracellular elicitin from *Phytophthora infestans*^57^), were transformed into *A. tumefaciens* by electroporation as described previously^22^.

### *Agrobacterium tumefaciens-mediated* transient transformation assays (ATTAs)

DsCEs were screened for their ability to elicit cell death in the non-host plants *N. tabacum* and *N. benthamiana* using an ATTA, as previously described^22,33^. Here, single colonies of *A. tumefaciens* transformed with a DsCE expression vector were first incubated in selective lysogeny broth (LB) at 28°C overnight. Cells were then collected by centrifugation at 3000× *g* for 5 min, resuspended in infiltration buffer (10 mM MgCl_2_, 10 mM MES (Sigma-Aldrich, St. Louis, MO, USA) in KOH, pH 5.6, 0.2 mM acetosyringone (Sigma-Aldrich, St. Louis, MO, EUA) to an OD_600_ of 0.5. Resuspended cultures were incubated at room temperature (RT) for at least 2 h, and infiltrated into the abaxial side of 5–6-week-old *N. tabacum* and *N. benthamiana* leaves. At least 12-24 infiltration zones were tested for each treatment (consisting of two leaves from each of two plants x 3-6 repeat experiments). In these ATTAs, INF1 was used as a positive control, while the empty pICH86988 vector was used as a negative control.

To verify the presence of DsCE proteins in agro-infiltrated *N. benthamiana* leaves, western blotting was carried out as described previously^39^. Here, total protein extracts from the leaves were separated by SDS-PAGE (12% bis-tris-acrylamide) and transferred onto PVDF membranes. Mouse anti-FLAG(R) antibodies (Sigma Aldrich) and chicken anti-mouse antibodies conjugated with horseradish peroxidase (Santa Cruz Biotechnology, Dallas, TX, USA) were used for detection by an Azure c600 imager (Azure Biosystems, Dublin, CA, USA) after incubation with SuperSignal^™^ West Dura Extended Duration Signal Substrate (Thermo Fisher Scientific, Waltham, MA, USA).

### Heterologous candidate effector protein expression in *P. pastoris* and protein purification

For the heterologous expression of Ds70057 in *P. pastoris*, the cDNA sequence encoding the protein, without its native signal peptide, was PCR-amplified from the previously generated ATTA expression vector. The cDNA sequence for BcSSP2 was amplified from the plasmid pPICZα-BcSSP2^37^. The primers used are shown in Supplementary Table S5. Resulting PCR amplicons from both candidates were cloned into the expression vector pPic9-His_6_ (Invitrogen, Carlsberg, CA, USA) behind the α-factor signal peptide sequence using *Sma*I/*Eco*RI restriction enzymes and T4 DNA ligase (New England Biolabs, Beverly, MA, USA). Here, the PCR 5’ primer was designed to incorporate a flag-tag for subsequent detection by western blotting. *P. pastoris* expression vectors were subsequently transformed into *E. coli*, and their sequence confirmed, as described above. Finally, the expression cassettes were linearized with *Sac*I or *Sal*I restriction enzymes (New England Biolabs) and transformed into *P. pastoris* according to Kombrink (2012)^58^.

Production of DsCE proteins in liquid culture using *P. pastoris* was performed according to Weidner et al. (2010)^59^. Here, expression of candidate proteins was induced by incubation of *P. pastoris* in 200 mL of BMMY (Buffered Methanol-complex Medium)^59^ for 72 h, with the successive addition of methanol every 24 h to increase the final concentration by 0.5% (v/v). Following incubation, *P. pastoris* cells were collected by centrifugation at 4500 g for 30 min. The supernatant, containing the secreted protein of interest, was sterilised through a 0.22 μm filter (ReliaPrep, Ahlstrom, Helsinki, Finland) and adjusted to pH 8 through the addition of NaOH.

Secreted candidate proteins were purified by immobilized metal ion affinity (IMAC) using Ni Sepharose 6 Fast Flow (GE Healthcare, Chicago, IL, EUA) according to the manufacturer’s protocol. Before loading, the column (Glass Econo-Column, BioRad, Hercules, CA, EUA) was packed with 5 mL of resin and was equilibrated by washing with binding buffer (20 mM sodium phosphate, 0.5 M NaCl, pH 7.4). The culture filtrate was added to the column and passed through at 1 mL/min. The protein was eluted with an elution buffer (20 mM sodium phosphate, 0.5 M NaCl and 500 mM imidazole, pH 7.4). The elution fractions were mixed, and elution buffer was added to a final volume of 50 mL to obtain enough volume to vacuum-infiltrate pine shoots. A western blot was performed, as above, to determine the presence of the proteins in *P. pastoris* culture filtrate and after purification.

### Pine infiltration with purified proteins

An overview of the pine infiltration method is shown in Fig. 3. Clonal rootless pine shoots were produced by adventitious shoot production from cotyledons^36^. The shoots were kept in LPch agar^36^ in glass jars, with each jar containing between six and eight shoots and three jars used for each treatment. For our experiment, two susceptible (S6 and S11) and one resistant (R4) genotype (based on field data) were used.

The shoots were completely submerged in approx. 50 mL of each purified DsCE or BcSSP2 protein solution, or in elution buffer or bovine serum albumin (BSA) (Thermo Fisher Scientific) as negative controls. Samples were exposed to vacuum in a glass chamber for 5 min, prior to gentle release of the vacuum to allow infiltration of the solution. Shoots were then rinsed in sterile MQ water, briefly air-dried, then placed back in the agar medium in the glass jar. The shoots were maintained in a 22°C room with natural light. Photos were taken 7 days after infiltration (dai) using a Nikon D7000 camera.

## Data Availability

The datasets analysed during the current study are available in the Joint Genome Institute and NCBI databases repository [https://genomes.jgi.doe.gov and https://www.ncbi.nlm.nih.gov/genbank]. All data generated during this study are included in this published article and its Supplementary Information files.

## Acknowledgements

This research was funded by Scion (NZFRI, Ltd.) through the Resilient Forests Research Program via Strategic Science Investments Funds from the New Zealand Ministry of Business Innovation and Employment (MBIE) and New Zealand Forest Grower Levy Trust funding. We thank the Kee Sohn Lab (formerly School of Agriculture and Environment, Massey University, New Zealand) for providing several *A. tumefaciens* strains and advice on the Golden Gate system. Paul Watson from the Radiata Pine Breeding Company is acknowledged for permissions to use pine material.

## Author contributions

RB, RM, MG and CM conceived and guided the study. LH and MT designed and performed experiments and analysed data. LH, MT and RB led manuscript writing. RM, CH and KG assisted with sourcing of pine materials and designing pine assays. CM and TL assisted with experiments. All authors contributed to the manuscript.

## Supplementary Figure Legends

**Figure S1: Locations of the 30 cloned *Dothistroma septosporum* candidate effector (DsCE) genes in the NZE10 genome.** The 14 chromosome-level scaffolds of *D. septosporum* NZE10 ^19^ are represented by the outer bars. Each minor tick represents 5,000 bp from the start of the scaffold; yellow stripes indicate the location of curated repetitive elements >200 bp in length ^32^. Outer numbers are protein IDs corresponding to the 30 DsCEs, with cell death-inducing DsCEs in bold font. For reference, the positions of the dothistromin biosynthesis genes^19^ are also shown (chromosome 12, grey labels). Within the inner rings, grey bars represent the 875 genes encoding putatively secreted proteins, and green bars (innermost) represent the 397 *in planta*-expressed (>50 Reads Per Million per Kilobase) secreted proteins. The figure was created using CIRCOS (http://circos.ca/software/^60^).

**Figure S2: Optical density (OD) range trial of *Agrobacterium tumefaciens* cultures.** Concentration thresholds are indicated for cell death elicitation by DsEcp2-1, but not Ds70057 and Ds131885. The yellow numbers show the used culture OD_600_; the positive (INF1) and negative (EV) controls were infiltrated at an OD_600_ of 0.6. In the Ds131885 panel, INF1 was trialled at 0.05 to 0.6 (top to bottom).

**Figure S3**: **Western blots of *Dothistroma septosporum* candidate effector proteins.** Western blots showed that DsCEs triggering cell death in *Nicotiana benthamiana* were expressed in the plant tissue regardless of the presence of a secretion signal peptide and absence of a cell death response (constructs with deleted signal peptide sequences are shown). Immuno-detection was based on primary anti-FLAG antibody.

**Figure S4**: **Replicates of *Pinus radiata* shoot tissue infiltrated with candidate effector proteins.** BcSSP2 (a) and Ds70057 (b) were produced by heterologous expression in *Pichia pastoris*. Photos were taken 7 days after infiltration.

