## Supplemental Figures for "Apoplastic effector candidates of a foliar forest pathogen trigger cell death in host and non-host plants"

### Hunziker et al. Supplementary Figures S1-S4

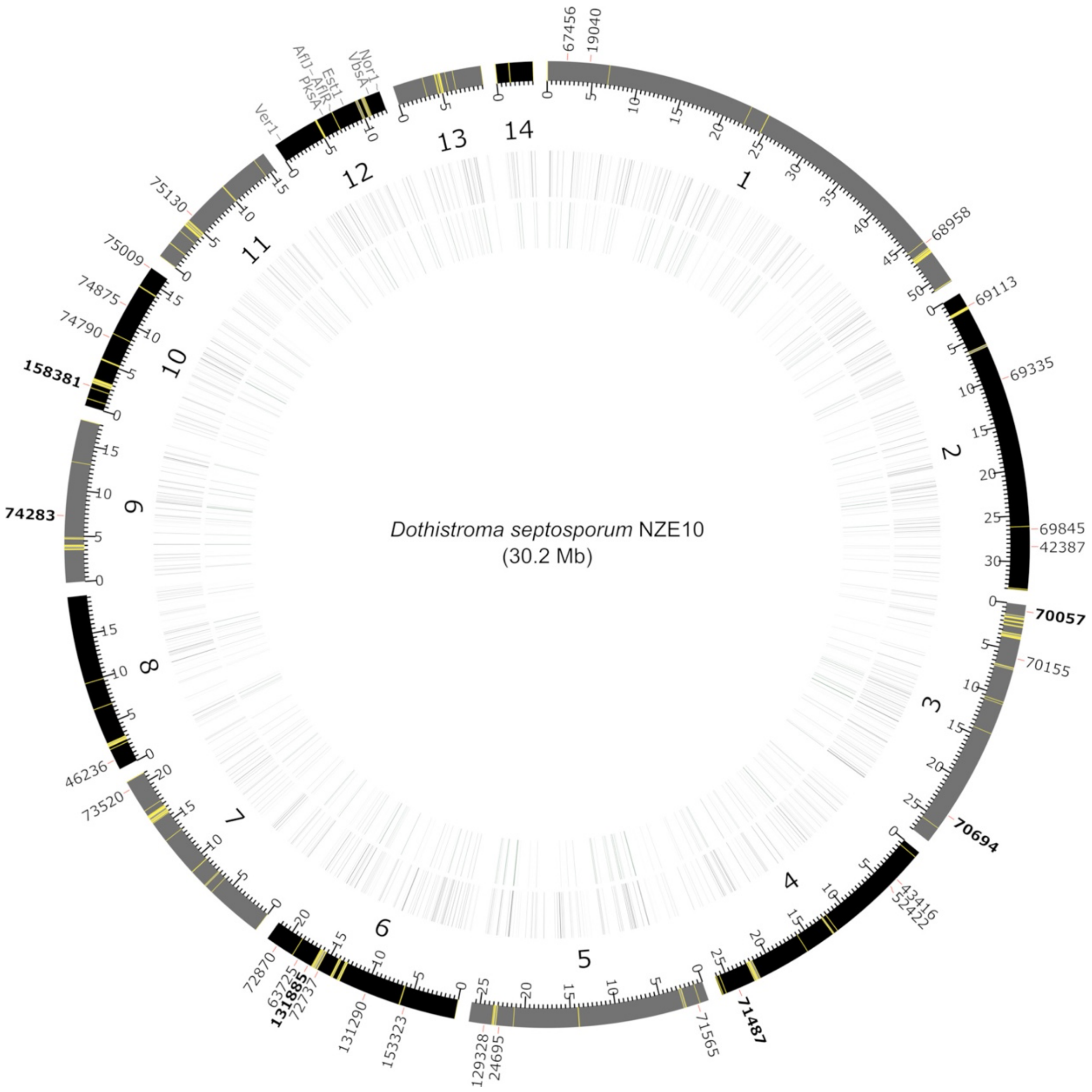


**Figure S1: Locations of the 30 cloned *Dothistroma septosporum* candidate effector (DsCE) genes in the NZE10 genome.** The 14 chromosome-level scaffolds of *D. septosporum* NZE10 [1] are represented by the outer bars. Each minor tick represents 5,000 bp from the start of the scaffold; yellow stripes indicate the location of curated repetitive elements >200 bp in length [2]. Outer numbers are protein IDs corresponding to the 30 DsCEs, with cell death inducing DsCEs in bold font. For reference, the positions of the dothistromin biosynthesis genes [1] are also shown (chromosome 12, grey labels). Within the inner rings, grey bars represent the 875 genes encoding putatively secreted proteins, and green bars (innermost) represent the 397 *in planta*-expressed (>50 Reads Per Million per Kilobase) secreted proteins. The figure was created using CIRCOS (http://circos.ca/software/; [3]).

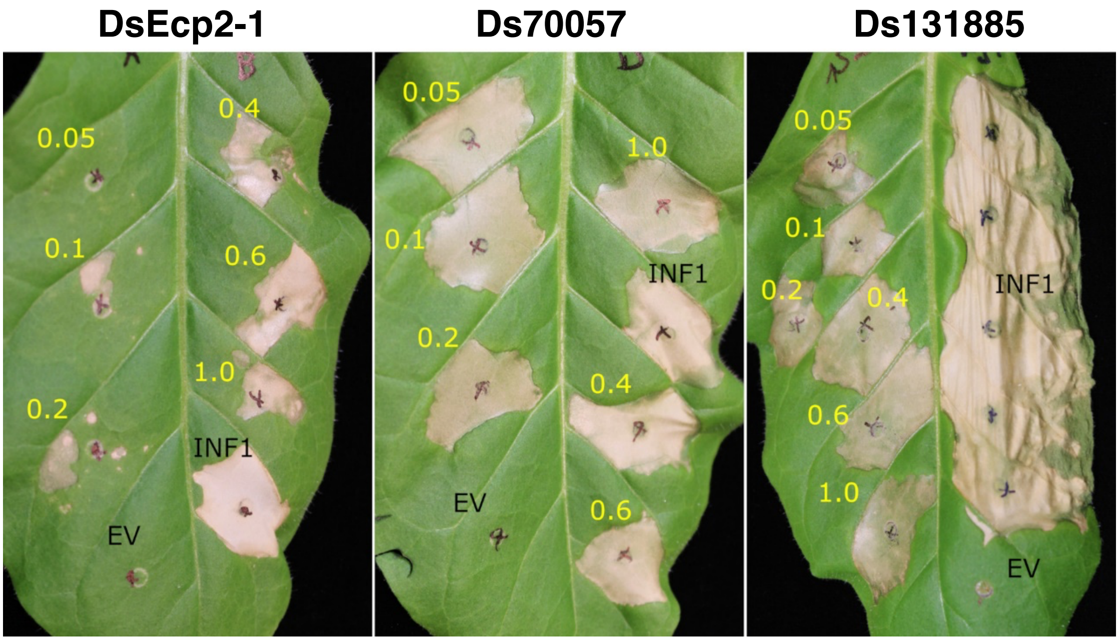


**Figure S2: Optical density range trial of *Agrobacterium tumefaciens* cultures.** Concentration thresholds are indicated for cell death triggering by DsEcp2-1, but not Ds70057 and Ds131885. The yellow numbers show the used culture $OD_{600}$; the positive (INF1) and negative (EV) controls were infiltrated at an $OD_{600}$ of 0.6. In the Ds131885 panel, INF1 was trialled at 0.05 to 0.6 (top to bottom).

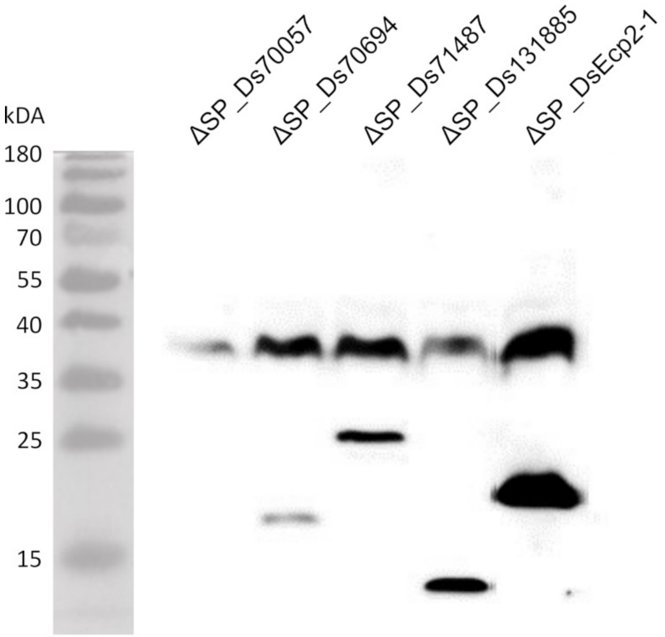


**Figure S3**: **Western blots of *Dothistroma septosporum* candidate effector proteins.** Western blots showed that DsCEs triggering cell death in *Nicotiana benthamiana* were expressed in the plant tissue regardless of the presence of a secretion signal peptide and absence of a cell death response (constructs with deleted signal peptide sequences are shown). Immuno-detection was based on primary anti-FLAG antibody.

(a)

BcSSP2 purified (1.3 µg/mL)

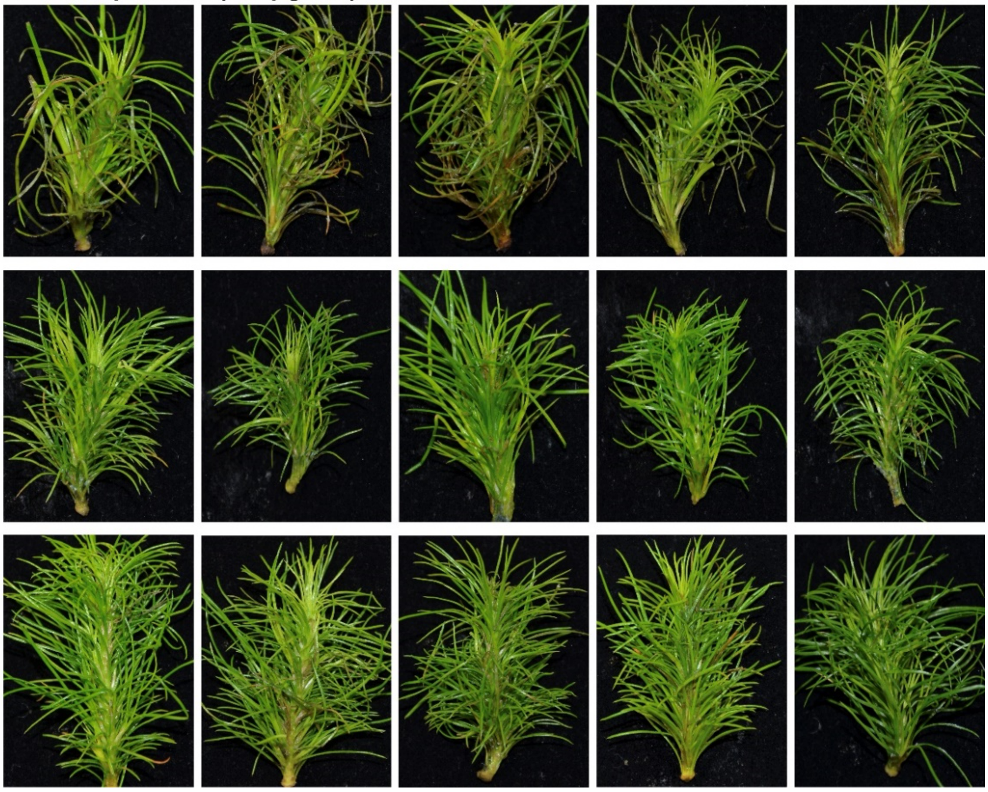


(b)

Ds70057 purified (21 µg/mL)

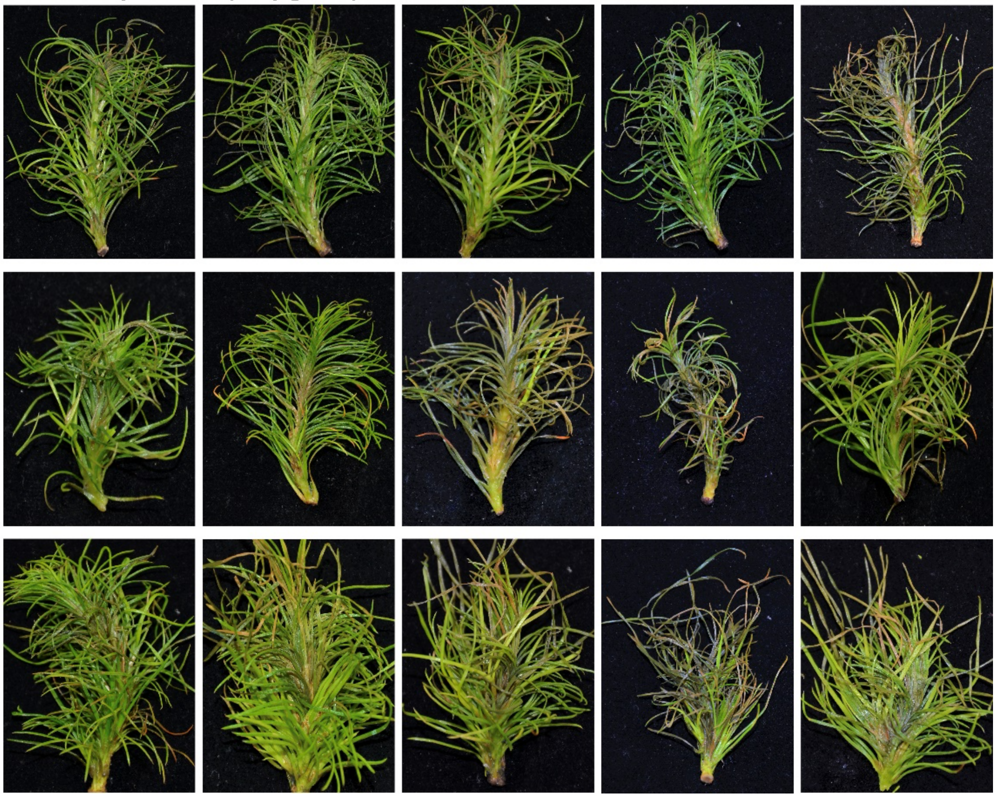


**Figure S4**: **Replicates of *Pinus radiata* shoot tissue infiltrated with candidate effector proteins.** BcSSP2 (a) and Ds70057 (b) were produced by heterologous expression in *Pichia pastoris*. Photos were taken 7 days after infiltration.
